# Accelerating single-cell genomic analysis with GPUs

**DOI:** 10.1101/2022.05.26.493607

**Authors:** Corey Nolet, Avantika Lal, Rajesh Ilango, Taurean Dyer, Rajiv Movva, John Zedlewski, Johnny Israeli

## Abstract

Single-cell genomic technologies are rapidly improving our understanding of cellular heterogeneity in biological systems. In recent years, technological and computational improvements have continuously increased the scale of single-cell experiments, and now allow for millions of cells to be analyzed in a single experiment. However, existing software tools for single-cell analysis do not scale well to such large datasets. RAPIDS is an open-source suite of Python libraries that use GPU computing to accelerate data science workflows. Here, we report the use of RAPIDS and GPU computing to accelerate single-cell genomic analysis workflows and present open-source examples that can be reused by the community.

## Introduction

Single-cell genomic technologies allow the profiling of diverse genome-scale information, including genome sequence, gene expression, chromatin accessibility, DNA methylation, protein-DNA binding and histone modifications, at the resolution of individual cells. Many computational tools have been created to analyze different types of single-cell genomic datasets. Two common frameworks that incorporate multiple analyses for single-cell RNA sequencing (scRNA-seq) data are Seurat [1] in R, and Scanpy in Python, which previously demonstrated speedups of 5 to 90 times relative to Seurat depending on the analysis step [2]. Similar frameworks to analyze single-cell ATAC-seq (scATAC-seq) data have been developed in R [3,4] and are being developed in Python [5].

Recently, new experimental methods and technological improvements have increased the number of single cells that can be profiled in a single experiment [6–8]. At the same time, improved algorithms for batch correction and data integration have enabled the construction of larger datasets by effectively integrating results from multiple experimental studies [9,10]. The increase in the number of cells analyzed per experiment translates into more data points, enabling increased power to distinguish between cellular states, but also requiring computational tools to scale rapidly. At the same time, there is a demand for tools that allow interactive, exploratory data visualization and analysis, necessitating a lower latency in results. Consequently, scaling to larger datasets has been described as a “sweeping theme” of single-cell biology [11]. A recent analysis of trends in the single-cell field also suggested that the increased scale of data generation has spurred development of methods to integrate datasets and classify cells using large reference datasets [12].

Here we report the use of RAPIDS and GPU computing to accelerate the analysis and visualization of single-cell genomic data, focusing on scRNA-seq and scATAC-seq data. RAPIDS is a suite of open-source Python libraries from NVIDIA that use Graphics Processing Unit (GPU) computing to accelerate data science workflows [13,14]. RAPIDS uses CUDA primitives for low-level compute optimization, and exposes GPU parallelism and high-bandwidth memory speed through Python interfaces. Our workflow uses several RAPIDS libraries: cuDF for dataframe manipulation, cuML for machine learning algorithms, and cuGraph for accelerated graph analytics.

We developed a series of examples demonstrating RAPIDS and GPU computing for different use cases in single-cell genomics, which are freely available in the form of Jupyter notebooks at https://github.com/clara-parabricks/rapids-single-cell-examples and are easily reusable. Using these, we obtained dramatic acceleration of single-cell workflows, particularly in key steps such as clustering and UMAP visualization. We show that GPU acceleration enables real-time, interactive analyses even on very large datasets containing over 1 million cells.

## Results

### RAPIDS accelerates single-cell RNA-seq analysis

We first accelerated a common scRNA-seq analysis workflow based on tutorials provided by Seurat and Scanpy [15,16]. Beginning with the scRNA-seq count matrix, we performed preprocessing (consisting of filtering cells and genes, normalizing the count matrix, subsetting the dataset to highly variable genes, regressing out confounding factors, and converting gene expression values to z-scores), dimension reduction, visualization, cell type identification via clustering, and differential gene expression analysis, using RAPIDS libraries throughout (Figure 1).

**Figure 1.**
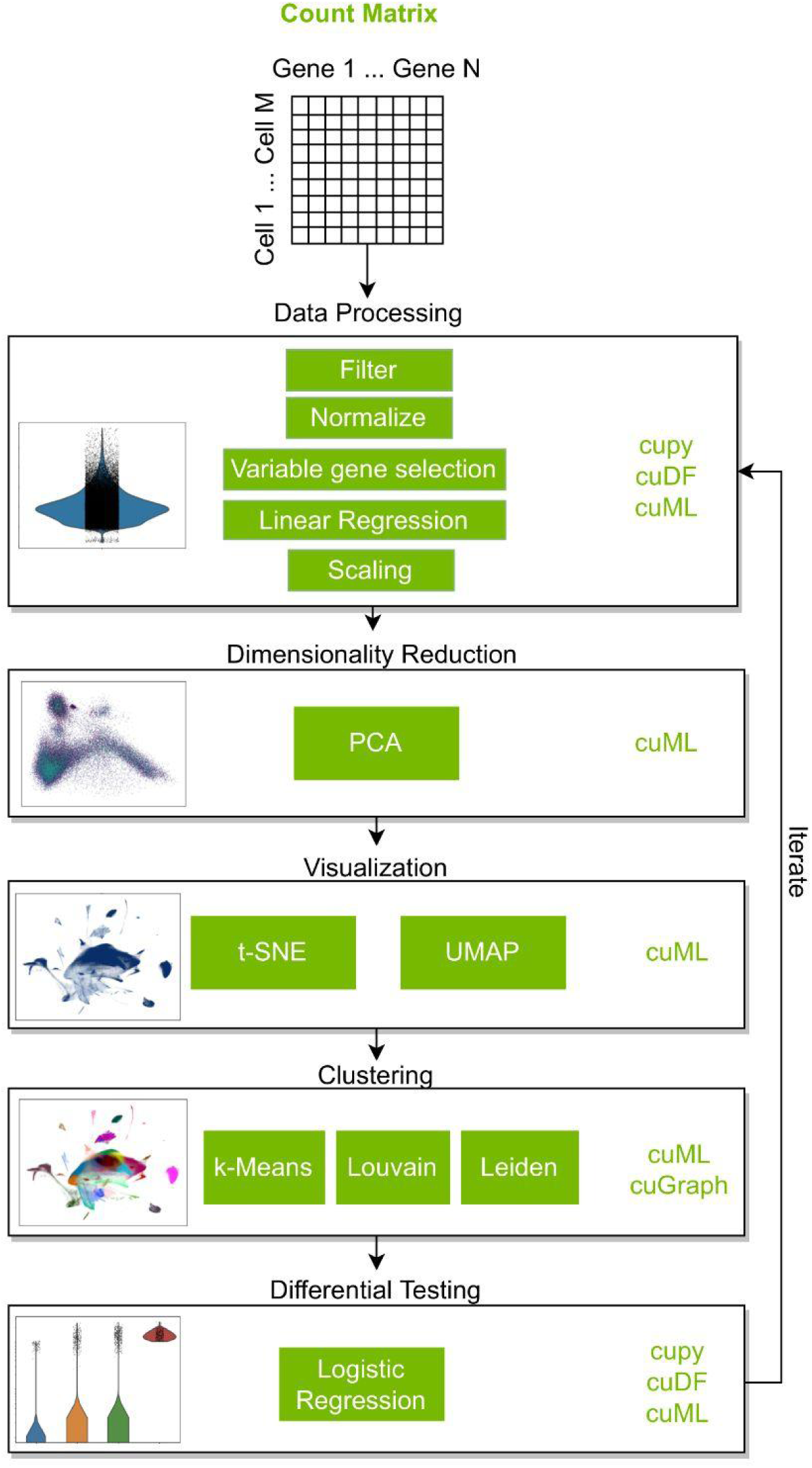
Schematic of the single-cell RNA-seq analysis workflow accelerated using RAPIDS libraries. A count matrix containing the expression values for each gene in each cell is preprocessed by filtering cells and genes, normalization, selecting highly variable genes, regressing out the effects of total counts and mitochondrial gene expression, and finally converting the values for each gene to z-scores. The processed count matrix is then reduced using PCA, and visualized using t-SNE and UMAP. The cells are clustered using k-means clustering, Louvain clustering, and Leiden clustering. Logistic regression is used to identify genes whose expression is significantly associated with cluster identity.

**Figure 2:**
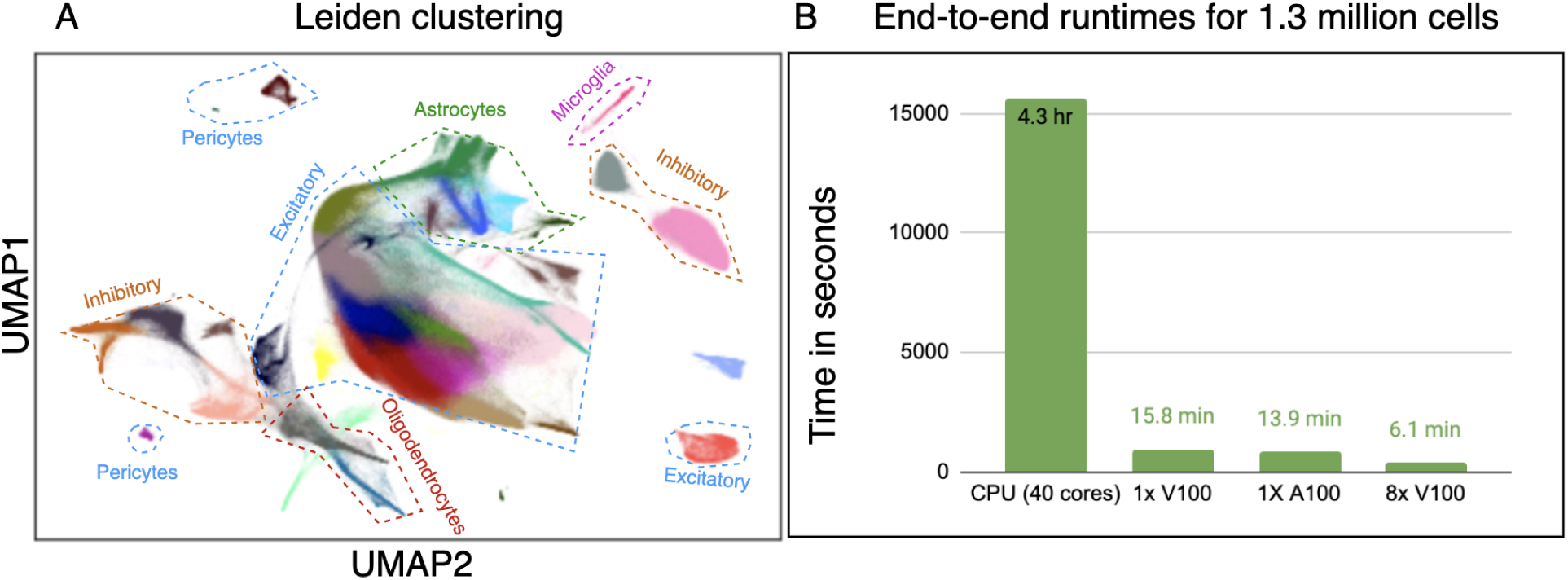
A. UMAP visualization of 1.3 million mouse brain cells labeled with Leiden clustering. Analysis was performed using a single V100 GPU. Major cell type labels were assigned based on marker gene expression (Figure S1). B. Comparison of runtime in seconds for a typical single-cell RNA workflow on 1.3 million mouse brain cells with different hardware configurations (Data in Table 1 and Table S1).

This was made easier by the fact that a few components of the RAPIDS workflow - specifically, RAPIDS K-Nearest Neighbors (KNN) graph construction, UMAP visualization, and Louvain clustering, had previously been integrated into the Scanpy framework [2]. In order to accelerate the complete workflow, we additionally implemented new functions for preprocessing and logistic regression on single-cell count matrices using cupy, cuDF and cuML operations, and used existing cuML and cuGraph functions to perform PCA, t-SNE, k-means clustering and Leiden clustering.

We stored our single-cell count matrix as an AnnData object [2] in order to take advantage of the data annotation, read/write functions, and visualization capabilities of the Scanpy/AnnData framework, as well as the RAPIDS algorithms already integrated into Scanpy. It should be noted that this is not the most time-efficient method, as the AnnData object is stored on the CPU, necessitating data transfer from the GPU to the CPU for storage in the AnnData object after every computation. Nevertheless, we structured our workflow in this way to optimize ease of use.

Single-cell count matrices are very sparse; for example, >90% of the elements in several of the datasets we tested were zeros. Making efficient use of memory required sparse formats and the ability to manipulate them easily required several additions to the CuPy library in order to provide features needed for preprocessing count matrices on the GPU, such as the ability to index and slice sparse arrays as well as perform reductions and binary arithmetic between them. Addition of the new sparse features in CuPy provided the additional benefit of enabling the Dask library to directly support distributed CuPy arrays.

Facebook’s FAISS library [17] provides a brute-force KNN which is used in RAPIDS cuML, along with some novel optimizations, by many of the neighborhood-based methods. The RAPIDS cuML library also contains novel GPU-accelerated implementations of UMAP [18], k-means clustering, linear, and logistic regression while the cuGraph library contains novel GPU-accelerated implementations of Louvain and Leiden clustering. cuML also contains GPU-accelerated Barnes-Hut [19] and FFT-interpolated [20] t-SNE variants, ported from the CannyLabs tsne-cuda library (https://github.com/CannyLab/tsne-cuda) to use RAPIDS C++ primitives. Further, RAPIDS makes heavy use of the Dask library to scale workloads across multiple GPUs and multiple physical machines [13]. Aside from being able to process data in sparse formats at scale, all of the cuML and cuGraph algorithms outlined in this paper are distributed in Dask, with the exception of t-SNE.

We scaled this workflow to 1.3 million cells using a dataset of mouse brain cells from 10X Genomics [21]. By sampling different numbers of cells from the 1.3 million cell dataset, we measured how the runtime of CPU and GPU algorithms scales with dataset size. All steps were executed on an NVIDIA DGX-1, which contains 8xV100 GPUs and Dual 20-core Intel Xeon processors. Although preprocessing the whole dataset at once on a single GPU was not possible for the complete 1.3 million cell dataset, it could be performed in batches, returning the same results. Another issue, currently unresolved, is that the t-SNE implementation in cuML is not fully reproducible even after fixing the random state.

The complete analysis of all 1.3 million cells was prohibitively slow on CPU, taking over 4 hrs. However, we observed significant acceleration in our GPU accelerated single-cell workflows, particularly for t-SNE, k-means, UMAP, Louvain clustering and Leiden clustering (Table 1). We further adapted the GPU workflow to use multiple GPUs and measured how the runtime of different steps scaled with the number of GPUs (Table 1). The complete analysis time was reduced to ∼16 minutes using a single GPU and further to just over 6 minutes using all 8 GPUs in parallel.

**Table 1.** scRNA-seq runtime for various steps using either 40 CPU cores on an NVIDIA DGX-1, or 1-8 V100 GPUs on a DGX-1. The ‘filtering and normalization’ step includes cell filtering, gene filtering, count matrix normalization, log transformation and selection of highly variable genes. All times are reported in seconds. Corresponding benchmarks for A100 GPU cloud instances are given in Table S1.

|  |  | CPU<br>40<br>cores | 1x<br>V100 | CPU<br>40<br>cores | 1x<br>V100 | CPU<br>40<br>cores | 1x<br>V100 | 2x<br>V100 | 4x<br>V100 | 8x<br>V100 |
| --- | --- | --- | --- | --- | --- | --- | --- | --- | --- | --- |
|  | Number of cells | 50,000 |  | 250,000 |  | 1.3M |  |  |  |  |
| Analysis Stage | Step |  |  |  |  |  |  |  |  |  |
| Preprocessing | Filtering & Normalization | 16.12 | 16.7 | 77.04 | 55.2 | 509.4 | 272 | 136.4 | 83.2 | 60.47 |
|  | Regression | 23.3 | 19.6 | 91.27 | 49 | 535.3 | 78 | 87 | 41.9 | 40.9 |
|  | Scaling | 1.65 | 1.21 | 8.51 | 0.842 | 52.1 | 42.4 | 14.2 | 4.63 | 2.39 |
| Dimensionality reduction | PCA | 7.63 | 1.13 | 37.5 | 18.6 | 199 | 152 | 81 | 35 | 11.4 |
| Visualization | t-SNE | 118 | 1.36 | 862 | 5.41 | 6093 | 52.1 | 52.5 | 52.3 | 52.3 |
|  | KNN | 16.2 | 6.76 | 32.3 | 13.6 | 142 | 87 | 88 | 89 | 91 |
|  | UMAP | 48.6 | 0.287 | 290 | 2.59 | 2234 | 57 | 57.9 | 57.8 | 57.8 |
| Clustering | K-means | 17.4 | 0.207 | 70 | 0.428 | 268 | 2.09 | 1.38 | 1.35 | 1.31 |
|  | Louvain | 8.25 | 0.116 | 87 | 0.558 | 707 | 3.04 | 3.38 | 3.01 | 3.1 |
|  | Leiden | 11.7 | 0.0902 | 201 | 0.476 | 3634 | 2.31 | 2.69 | 2.24 | 2.37 |
| Differential gene expression |  | 52.4 | 3.28 | 221 | 20.8 | 1155 | 149 | 3.39 | 3.26 | 3.94 |
| Full Workflow |  | 411.38 | 62.04 | 2075 | 184.9 | 15656 | 951.4 | 578.42 | 412.7 | 365.4 |

Comparing the results of the analysis on CPU and GPU analysis on the full 1.3 million cells, we found that the scaled and centered count matrices were nearly identical (fewer than .01% of the cells in the count matrix differed by more than 0.1% of their value), and the top 50 PCA components found by the two pipelines had an absolute Pearson correlation of at least 0.999. Further, the percentage of nearest neighbor cells conserved between the two analyses, averaged across all cells, was 95.66%.

Comparing the results of Leiden clustering, we found that the CPU workflow resulted in 37 clusters whereas the GPU workflow resulted in 34. The number of clusters found by this algorithm depends upon the initial state and varies with the selection of different random seeds. However, the measured concordance between cluster assignments was high (Adjusted Rand Index = 0.54, Normalized Mutual Information = 0.76). Further, we identified major brain cell types (excitatory neurons, inhibitory neurons, astrocytes, oligodendrocytes, microglia, and pericytes) based on their marker gene expression (Figure S1), and found that cells expressing different marker genes were both well separated by GPU UMAP visualization, and were partitioned into distinct clusters by Leiden clustering on GPU.

### Building interactive single-cell data visualizations with RAPIDS

The speed of GPU-accelerated analyses enables interactive analysis even of very large datasets. This is especially dramatic in visualization and clustering steps, which can be performed in seconds even for the complete dataset of 1.3 million cells (Table 1, Table S1). This enables in-browser, interactive visual data analysis at scale, a powerful capability that is currently missing in popular interactive cell browsers [22,23].

As a demonstration of this capability, we built an in-browser, interactive visualization of scRNA-seq data using the interactive dashboarding library Plotly Dash [24] in combination with RAPIDS. The interactive dashboard launches from a Jupyter notebook and displays a 2-D visualization of the single-cell dataset. Users can select groups of cells, visualize their properties, and perform GPU-accelerated dimension reduction, visualization and clustering by pointing and clicking in the browser. Loading a dataset of 70,000 cells into the dashboard on a DGX-1 (using a single V100 GPU), we observed that we could interactively select groups of upto 15,000 cells, perform GPU-accelerated PCA, UMAP and Leiden clustering, and render a new UMAP visualization of the selected cells, in a total of under 10 seconds. This speed allows for even complex analyses to be performed in a code-free, visual and interactive workspace without delays that break the user’s working flow.

### Accelerating scATAC-seq analysis and visualizing genomic coverage tracks

Finally, we extended our work to scATAC-seq analysis, which includes many of the same steps as scRNA-seq. However, scATAC-seq provides genome-wide measurements that are commonly represented as a count matrix by calculating the per-cell coverage within either peaks or genomic bins, rather than genes. Thus the scATAC-seq count matrix commonly has more features than examples.

scATAC workflows are currently underdeveloped in Python, while more advanced frameworks have been developed for R [3,4]. We therefore developed our own python workflow for single-cell ATAC-seq analysis, based on the Signac tutorial in R [25]. We ran this analysis on a dataset of approximately 60,000 human PBMCs sequenced using the dsci-ATAC-seq protocol [26].

We found that preprocessing steps (specifically, peak and cell filtering, and TF-IDF normalization) were very fast on CPU, rendering GPU acceleration unnecessary for the number of cells in this dataset. However, the subsequent steps were accelerated using RAPIDS as for scRNA-seq, resulting in similar acceleration to that observed in scRNA-seq (Table 2). Notably, the acceleration of PCA was lower than that observed for scRNA-seq datasets with similar numbers of cells, due to the larger number of features relative to cells in scATAC data.

**Table 2:**
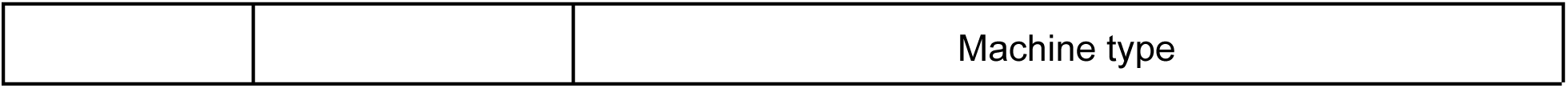
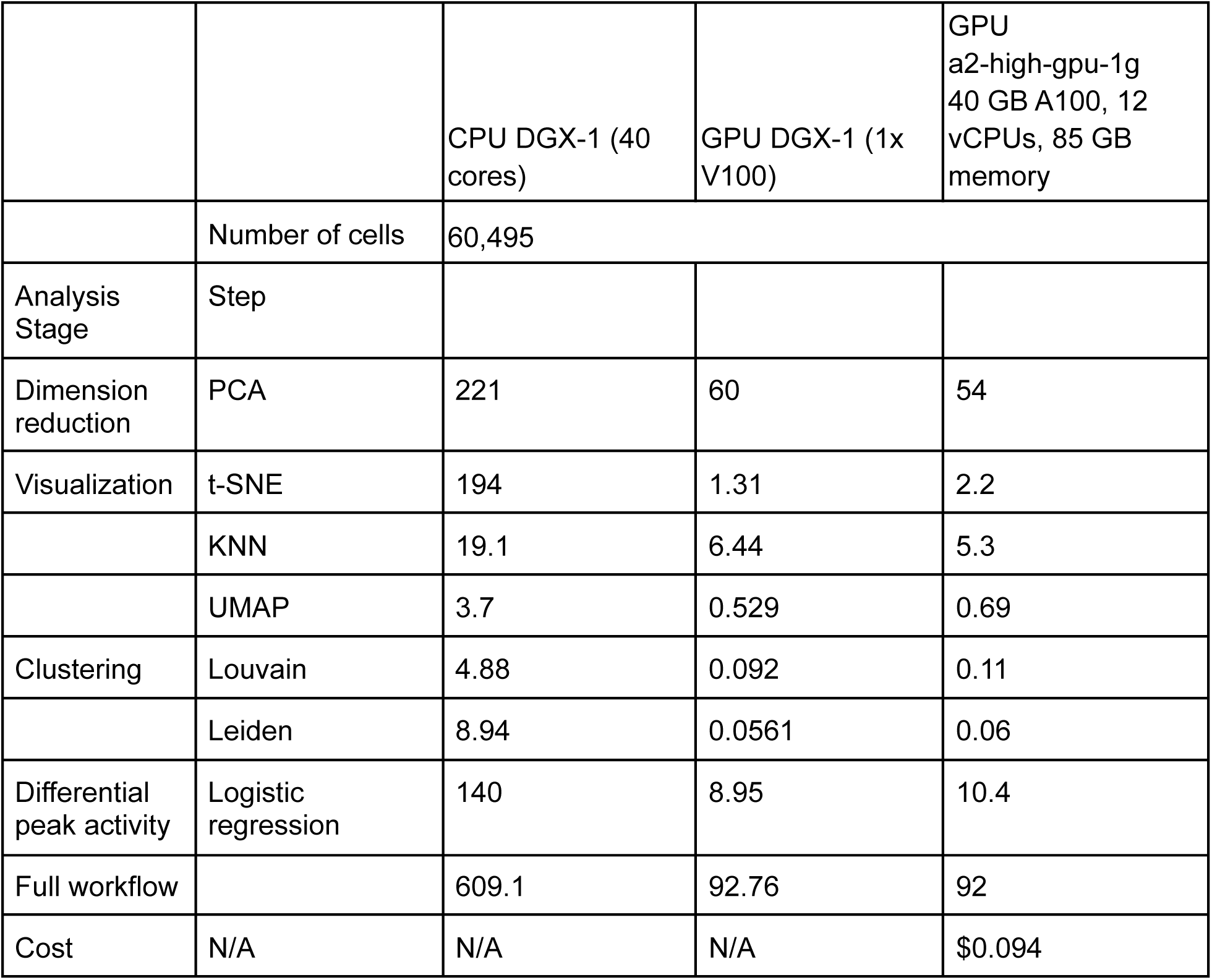
scATAC-seq runtime for various steps using either 40 CPU cores on an NVIDIA DGX-1, a single V100 GPU on a DGX-1, or a Google Cloud Platform (GCP) instance with a single A100 GPU. All times are reported in seconds.

|  |  |  |
| --- | --- | --- |
|  |  | Machine type |

|  |  |  |  |  |
| --- | --- | --- | --- | --- |
|  |  | CPU DGX-1 (40 cores) | GPU DGX-1 (1x V100) | GPU a2-high-gpu-1g 40 GB A100, 12 vCPUs, 85 GB memory |
|  | Number of cells | 60,495 |  |  |
| Analysis Stage | Step |  |  |  |
| Dimension reduction | PCA | 221 | 60 | 54 |
| Visualization | t-SNE | 194 | 1.31 | 2.2 |
|  | KNN | 19.1 | 6.44 | 5.3 |
|  | UMAP | 3.7 | 0.529 | 0.69 |
| Clustering | Louvain | 4.88 | 0.092 | 0.11 |
|  | Leiden | 8.94 | 0.0561 | 0.06 |
| Differential peak activity | Logistic regression | 140 | 8.95 | 10.4 |
| Full workflow |  | 609.1 | 92.76 | 92 |
| Cost | N/A | N/A | N/A | \$0.094 |

An additional challenge in interactive analysis for scATAC-seq data is the need to visualize local sequencing coverage around genes or regulatory elements of interest, such as cell-type specific enhancers and promoter regions, and compare sequencing coverage in such regions between clusters. However, it is a slow and limiting process to generate coverage tracks for each newly discovered cluster and visualize them in a genome browser.

We demonstrated how, following cluster identification, RAPIDS can be used to load an scATAC-seq fragment file using the cuDF library and create sequencing coverage tracks for selected regions in each cluster, thus enabling interactive cluster-specific visualization alongside interactive clustering. We further integrated this functionality with AtacWorks [27], a deep learning tool to enhance ATAC-seq coverage tracks and call peaks accurately even from low cell count data. This allowed us to not only rapidly identify and visualize clusters with GPU-accelerated algorithms, but also to generate real-time, in-notebook, deep-learning enhanced visualizations of sequencing coverage and peak calls in each cluster.

## Discussion

Here, we demonstrate the use of RAPIDS to accelerate single-cell genomic data analysis through a series of open-source examples. RAPIDS shows dramatic acceleration of limiting steps in current single-cell analysis workflows, such as clustering and visualization, and scales to massive datasets. As a demonstration, we analyzed scRNA-seq data from 1.3 million mouse brain cells and showed that the analysis time could be reduced from 4.3 hours to 6 minutes. This acceleration permits real-time, interactive analysis and visualization of very large datasets, potentially enabling a new family of GPU-accelerated cell browsers and other interactive visualization tools. We further demonstrated that this workflow can be adapted to obtain similar acceleration in the analysis and visualization of datasets from different genomic modalities, such as scATAC-seq. Adding GPU compatibility to the AnnData library and Scanpy would speed up these workflows further by reducing data transfer between the CPU and GPU.

Finally, new data-intensive challenges are presented by the growing list of multimodal single-cell methods which measure multiple high-dimensional genomic modalities from the same cell [28], thus increasing data dimensionality as well as the number of cells. At the same time, new algorithms are being developed to integrate datasets from different sources and modalities with batch correction [9,10], further increasing the number of cells to be analyzed at a time, and underscoring the importance of accelerated computing in this area. These trends, along with the acceleration observed in this study, suggest that the use of accelerated computing to enable faster, interactive analysis at scale will have a dramatic impact on single-cell genomic research.

## Data availability

The 1.3 million cell scRNA-seq dataset used here is available from 10X Genomics at https://support.10xgenomics.com/single-cell-gene-expression/datasets/1.3.0/1M_neurons. The scATAC-seq dataset used here was made public by Lareau et al. [26]. We processed the dataset to include only cells in the ‘Resting’ condition and peaks with nonzero coverage in at least one cell. Our processed files are available at https://rapids-single-cell-examples.s3.us-east-2.amazonaws.com/.

## Code availability

Our workflows are freely available under an Apache 2.0 license, along with a docker image containing all requirements, at https://github.com/clara-parabricks/rapids-single-cell-examples. The benchmarks reported in this manuscript were performed using release 22.02 which uses RAPIDS version 22.02 and Scanpy version 1.9.1.

