## Supplementary Material for "Accelerating single-cell genomic analysis with GPUs"

| | | a2-highgpu-1g | | a2-highgpu-2g | | a2-highgpu-4g | a2-highgpu-8g |
|---|---|---|---|---|---|---|---|
| | Number of cells | 50,000 | 250,000 | 1,300,000 | | | |
| Analysis Stage | Step | | | | | | |
| Pre-processing | Filtering & normalization | 110 | 158 | 395 | 138* | 83* | 57.48* |
| | Regression | 49.7 | 79 | 106 | 86* | 46.4* | 46.5* |
| | Scaling | 0.164 | 1.93 | 5.33 | 7.12* | 3.23* | 1.67* |
| Dimension reduction | PCA | 1.04 | 6.12 | 27.2 | 13.09* | 11.23* | 12.24* |
| Visualization | t-SNE | 2.01 | 6.52 | 36.7 | 37.5 | 37.4 | 37.1 |
| | KNN | 7.84 | 14.6 | 68 | 65 | 67s | 65s |
| | UMAP | 0.418 | 1.78 | 21.3 | 7.78* | 5.07* | 3.36* |
| Clustering | k-means | 0.130 | 0.442 | 2.12 | 5.84* | 5.07* | 8.44* |
| | Louvain | 0.146 | 0.535 | 2.75 | 2.72 | 2.67 | 2.82 |
| | Leiden | 0.121 | 0.377 | 1.92 | 1.97 | 1.9 | 2.06 |
| Differential gene expression | Logistic regression | 2.75 | 15.4 | 104 | 3.16 | 2.01 | 2 |
| Full workflow | | 184.84 | 304.95 | 835 | 401 | 290 | 271 |
| Cost | | $0.15 | $0.25 | $1.36 (1gpu) | $0.65 (2gpu) | $0.94 | $1.77 |

* parallelized over multiple GPUs
VM pricing: 2.939 per GPU per hour

***Table S1**: scRNA-seq runtime for various steps on GCP Cloud Instances. All steps were run on a Google Cloud Platform (GCP) instance with A100 GPUs (a2-highgpu-1g, a2-highgpu-4g, a2-highgpu-8g), and 12, 48, and 96 vCPUs respectively. All times are reported in seconds.*

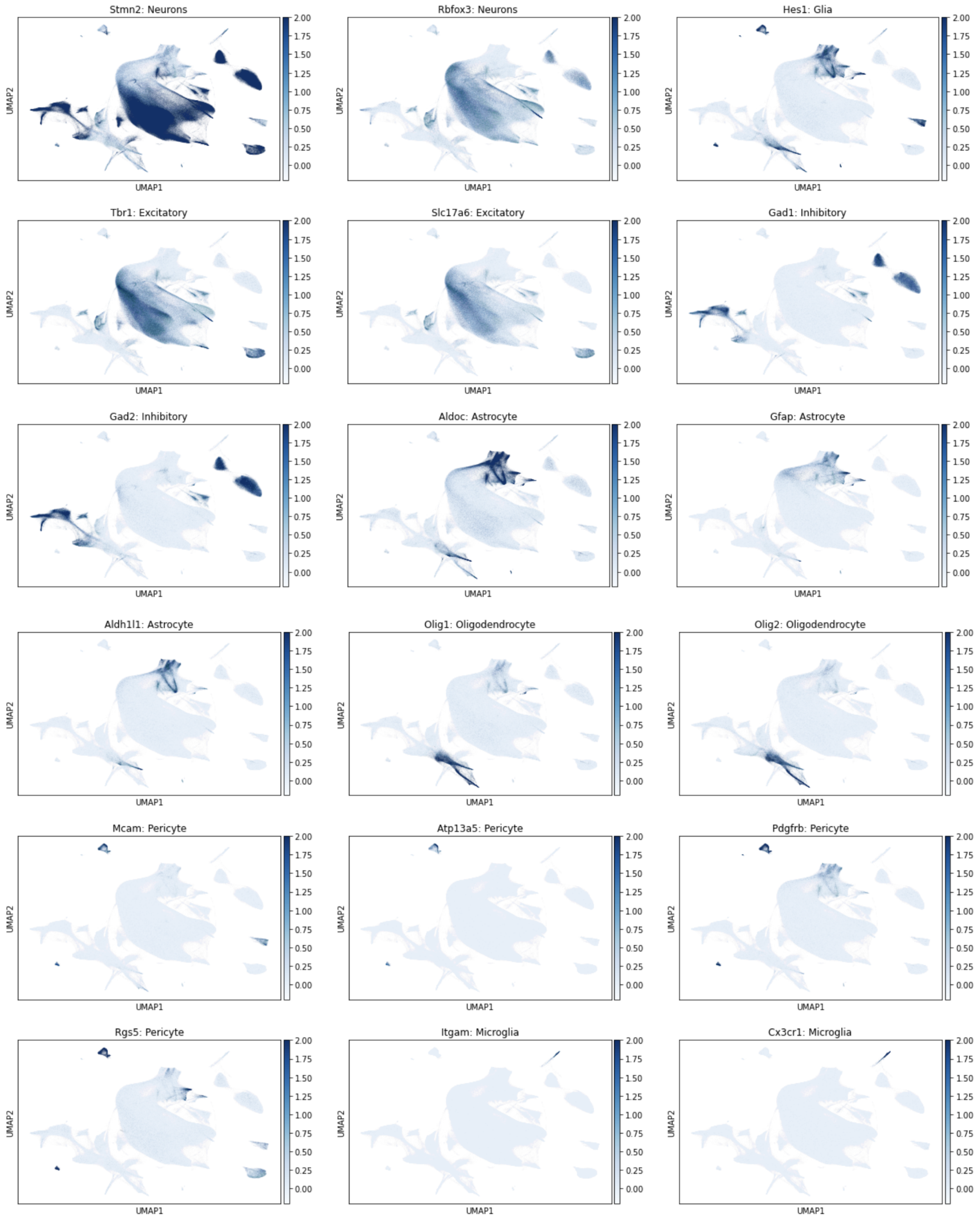

Stmn2: Neurons
Rbfox3: Neurons
Hes1: Glia
Tbr1: Excitatory
Slc17a6: Excitatory
Gad1: Inhibitory
Gad2: Inhibitory
Aldoc: Astrocyte
Gfap: Astrocyte
Aldh1l1: Astrocyte
Olig1: Oligodendrocyte
Olig2: Oligodendrocyte
Mcam: Pericyte
Atp13a5: Pericyte
Pdgfrb: Pericyte
Rgs5: Pericyte
Itgam: Microglia
Cx3cr1: Microglia
UMAP1
UMAP2
2.00
1.75
1.50
1.25
1.00
0.75
0.50
0.25
0.00

***Figure S1***: *UMAP visualization of expression of selected marker genes for major brain cell types, in the dataset of 1.3 million mouse brain cells from 10X genomics. The analysis was performed using a single V100 GPU. Values plotted are normalized and log-transformed counts. Values greater than 2 were set to 2 for visualization.*
